# Exploring the Impact of S. aureus Lipase Activity on Intra- and Extracellular Lipids

**DOI:** 10.64898/2026.07.29.741556

**Authors:** David T. Brewer, Kelly M. Hines

## Abstract

Previous research has shown that mammalian fatty acids (FAs) can influence antibiotic tolerance of *Staphylococcus aureus*, yet many of these studies overlook the sources of these FAs, which are primarily esterified into glycero- and phospholipids, and the impact *S. aureus* lipase activity has on host lipids. Here we attempt to gain insight into the complex interplay between the *S. aureus* lipidome and its environment using culture media supplemented tissue-specific phospholipid mixtures. Phospholipid profiles of heart, liver, and brain-derived lipids revealed distinct distributions of headgroup and fatty acyl tail structures within the phospholipids. Following the growth of *S. aureus* in lipid-enriched broth, PG species containing mono- and poly-unsaturated acyl tails were detected with abundances that correlated strongly with the FA profile of the tissue extract. We found that *S. aureus* cultured with liver-derived lipid extract, which yielded the most unsaturated PGs, promoted growth in high concentrations of the membrane-targeting antimicrobial daptomycin. To explore the influence of lipase activity on the extracellular lipids, comparative analysis of fresh versus spent media revealed that the lipase-mediated degradation of complex phospholipid mixtures was influenced by both head group structure and acyl tail linkage. Concurrently, the spent media contained elevated levels of mono- and polyunsaturated lysophospholipids that were predominantly of the 2-acyl form rather than the 1-acyl form observed in the fresh media. Together, these results demonstrate the extent to which the lipase activity of *S. aureus* remodels both its own lipidome as well as the structures of the phospholipids in the surrounding environment.

**IMPORTANCE:** *S. aureus* releases a secreted glycerol ester hydrolase, Geh, into the extracellular environment, which enables the bacterium to generate free FA from glycerolipids, phospholipids, and cholesterol esters that are present in surrounding tissue of an infection. The liberated FAs can be incorporated into the phospholipids of *S. aureus*, thereby altering its membrane physiology with mono- and poly-unsaturated FAs it cannot otherwise synthesize. Simultaneously, the action of Geh on lipids in the host environment leads to higher levels of bioactive lysophospholipids that participate in mammalian signaling pathways. This work reveals the preferences of *S. aureus* Geh across phospholipids with different head group and acyl tail structures found within tissue-derived lipid extracts, as well as the fate of the liberated FAs within the staphylococcal membrane lipids. The impacts of these processes on both the host and bacterium have implications for the immune response to and antibiotic treatment of *S. aureus* infections.

## INTRODUCTION

Nearly 15% of global *S. aureus* infections are fatal despite the general decline in health care and community-associated *S. aureus* infection rates over the past 13 years.^1^ While most often associated with skin infections, invasive *Staphylococcus aureus* infections following surgery, medical device implantation, or severe bloodstream infections can lead to the infection of organs including the heart, liver, and kidney.^2, 3^ The complications of such infections can lead to organ injury or failure, stroke, or sepsis.^3^ *S. aureus* uses an array of virulence factors ranging from secreted toxins to the ability to form resilient biofilms that enable its invasion into and survival within organs and tissues. Two of these systems target critical barriers formed by mammalian and staphylococcal cell membranes by altering the structure and composition of their lipids (**Figure 1**).

**Figure 1.**
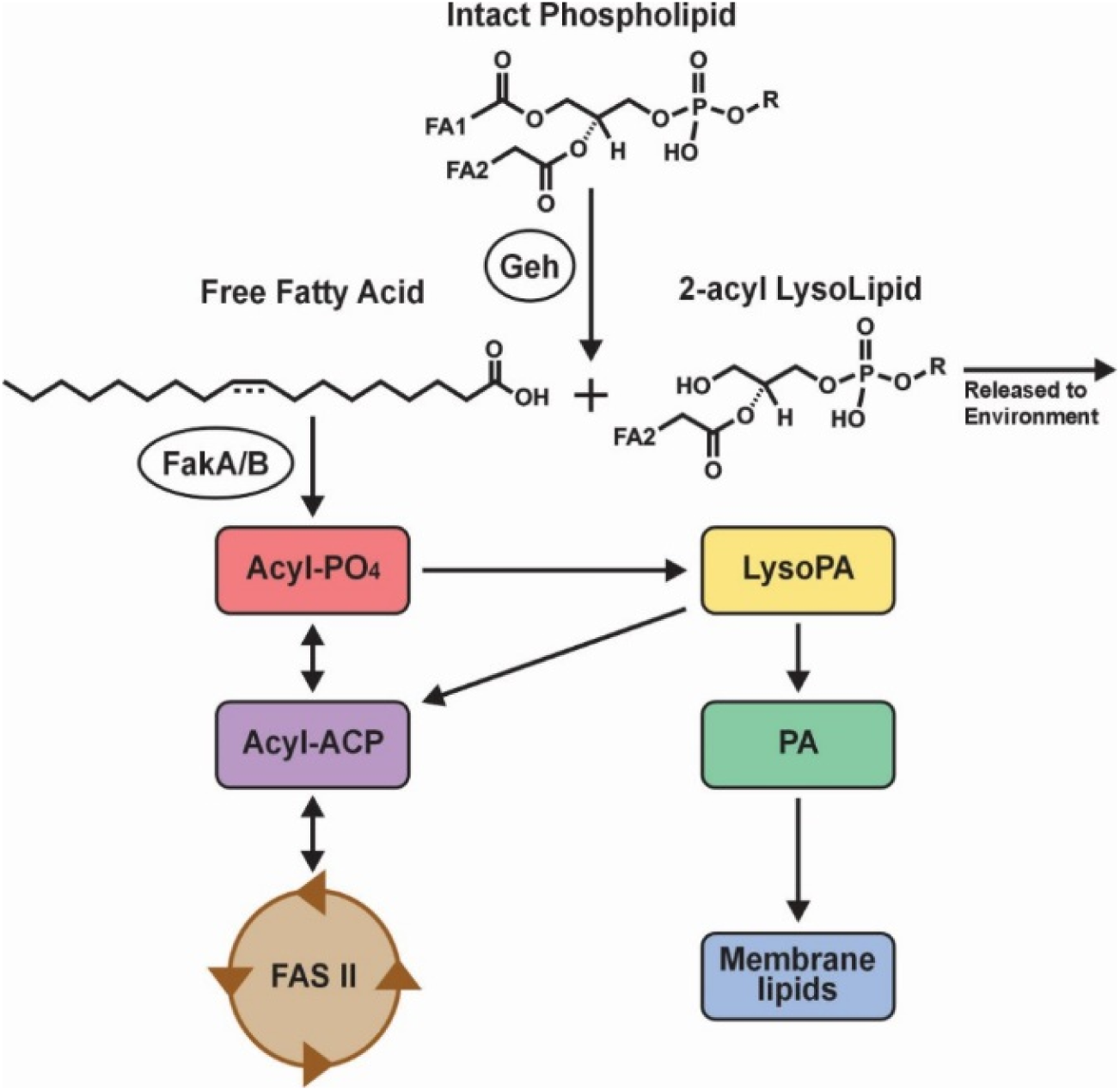
Glycerol ester hydrolase (Geh) and fatty acid kinase (Fak) pathways for the liberation and incorporation of exogenous free fatty acids.

The fatty acid kinase system (Fak) was discovered in 2014 as a mechanism to evade antimicrobials that were developed to target FASII.^4^ This system enables exogenous free fatty acids (FFAs) to enter the lipid biosynthetic pathway of *S. aureus* as an alternative to endogenous FFAs produced *de novo* by FASII. One of two fatty acid binding proteins, FakB1 or FakB2, position the FFA for phosphorylation into an acyl-phosphate by the fatty acid kinase, FakA.^4–6^ From this point, the exogenous FA can undergo elongation within FASII or enter the organism’s lipid biosynthesis pathways through a lysophosphatidic acid intermediate. The ability to use exogenous FAs in its membrane lipid synthesis presents several advantages for *S. aureus*. Firstly, *de novo* fatty acid synthesis is an energy-expensive process, and the ability to scavenge rather than synthesize FAs conserves energy in an environment that is challenging for *S. aureus* survival.^7^ Secondly, the direction of FAs into its own membrane lipids can serve as a detoxification mechanism to prevent damage from host-derived FAs with antimicrobial properties.^8–13^ Lastly, the Fak system provides *S. aureus* with unsaturated FA structures that it cannot produce itself, which can promote higher membrane fluidity and protect against membrane-targeting antimicrobials.^5, 14^

The amount of free fatty acids that are directly accessible for uptake through the Fak pathway is relatively small compared to other pools of fatty acids. Most FAs are esterified to the glycerol backbones of glycero- and glycerophospholipids, or to cholesterol to form cholesterol esters.^15–17^ To generate a source of FFAs, *S. aureus* can produce a secreted lipase, glycerol ester hydrolase (Geh, also known as SAL2 or Lip2), that cleaves ester linkages within triacylglycerols (TG), phospholipids (PL), and cholesterol esters (CE).^18^ Previous studies on Geh have shown that it can act on the lipids found within human serum and low-density lipoproteins to release FFAs, leaving diacylglycerols (DG), lysophospholipids (LysoPL), and free cholesterol.^18–21^ Further study of Geh has investigated the specific structural preferences of the lipase across different lipid species and positional isomers. Incubation of Geh with standards of the most abundant phospholipids in mammalian cells, phosphatidylcholine (PC) and phosphatidylethanolamine (PE), revealed that both classes of phospholipids can be acted upon by the lipase. Upon incubation of Geh with asymmetrical species of PC and PE, where palmitate at the *sn-1* position of the glycerol backbone and linoleate at the *sn-2* position, free linoleate was detected in the supernatant of PC 16:0/18:2 whereas linoleate did not rise above the backgrounds following incubation of Geh with PE 16:0/18:2.^22^ While these results were not quantitative and persistent background of FFAs complicated interpretation, the evidence of free linoleate in the supernatant suggests that Geh has a preference for either the *sn-2* position or unsaturated fatty acyl tails. However, incubations of Geh with phosphatidylglycerols (PGs) that make up the staphylococcal membrane suggests that proGeh favors the *sn-1* position of the backbone.^18^ Geh is also considered a virulence factor in *S. aureus* infections given its ability to inactivate components of innate immune defense. Recent work has demonstrated that glycerol released from the complete hydrolysis of triacylglycerols by Geh is used to promote biofilm formation in *S. aureus*.^23^ Despite evidence of Geh’s role in *S. aureus* pathogenicity, there is still much that is unknown regarding the interactions of Geh with the other classes and structures of phospholipids that are found in mammalian cells and tissues.

In this study, we evaluate the interaction of *S. aureus* with biologically relevant mixtures of mammalian phospholipids to reveal the impact of Fak and Geh activity on both the membrane lipids of *S. aureus* and the phospholipids in the extracellular domain. Lipid extracts derived from different organs were used to simulate the diversity of phospholipid classes and acyl tail compositions found across different mammalian tissues. By using lipid-enriched broths, we have reconstituted the lipid complexity of brain, heart, and liver tissue into *S. aureus* cultures. Our lipidomics results reveal that the *S. aureus* fatty acid profile mimics that of the lipid-enriched media, whereas the lipid composition of the spent media is significantly altered by the presence of *S. aureus*. This reductionist approach to replicating the infection environment provides new insight into the lipid content of the infection environment and the staphylococcal membrane *in vivo*, which have implications for host immune response and antimicrobial susceptibility of *S. aureus*.

## EXPERIMENTAL METHODS

### Materials and Chemicals

Heart, liver, and brain polar lipid extracts were obtained from Avanti Polar Lipids as powders and stored at 25 mg/mL in chloroform at -80°C. Tryptic soy broth, tryptic soy agar, ammonium acetate, ammonium formate, and HPLC grade solvents were purchased from Fisher Scientific. *Staphylococcus aureus* strain JE2 (NR-4653) was obtained from BEI Resources. Daptomycin was purchased from Gold Biotechnology and stored at 10 mg/mL in sterile water at -80°C.

### Bacteria and Growth Conditions

Bacteria were plated onto tryptic soy agar (TSA) and incubated overnight at 37°C. Colonies were then diluted to 2 McFarlands (∼ 6 x 10^8^ CFU/mL) using a benchtop densitometer (BEN-1B-Bio). Aliquots of heart, liver, or brain polar lipid extract were vacuum-dried (Savant, ThermoScientific) and reconstituted in DMSO to a concentration of 5 mg/mL. Tryptic soy broth (TSB) (10 mL) was spiked with 100 μL of individual lipid extract to a final concentration of 50 μg/mL, DMSO, or left untreated. Cultures (n=3) were prepared by adding 100 μL of bacterial suspension or vehicle control to 900 μL of TSB containing lipid extract or DMSO. Additional controls lacking bacteria were also prepared. The cultures were incubated overnight at 37°C with shaking at 200 rpm.

### Lipid Extraction

Lipid extractions were performed using a modified version of the biphasic Bligh and Dyer extraction.^24^ Following overnight growth, cultures were centrifuged for 10 minutes at 3000 rpm at 4 °C to pellet the bacteria. From each culture, 500 μL of spent media was transferred into 5 mL glass tubes for extraction, and the remaining broth was discarded. Bacterial pellets were washed twice and pelleted by centrifugation. Pellets were then suspended in 800 μL of HPLC water, from which 500 μL was used for lipid extraction. The remaining suspension was used to determine optical densities (OD600, BioTek Epoch 2). Bacteria suspensions were sonicated in an ice bath for 30 min, followed by the addition of 2 mL of chilled 1:2 chloroform/methanol (*v/v*) solution and intermittent vortexing for 5 min. Sequentially, 500 μL of chilled chloroform and 500 μL of HPLC water were added and vortexed for 1 min, and samples were centrifuged at 3000 rpm at 4 °C for 10 min to induce phase separation. The bottom organic layer was then transferred into new glass tubes, dried in a vacuum concentrator, and reconstituted with 0.5 mL of 1:1 chloroform/methanol (*v/v*) prior to storage at -80 °C.

### RPLC-IM-MS and -MS/MS Data Acquisition

Lipid extracts from bacteria pellets were dried under vacuum and reconstituted in 1:2 acetonitrile/methanol with a 2.5-fold dilution factor. The lipid extracts from broths were prepared similarly but diluted so that the concentration of tissue-derived lipids was 15 μg/mL. Quality control (QC) samples were generated by combining 10 μL of each bacteria or broth extract, respectively, into a single vial. Lipid extracts were analyzed using reverse-phase liquid chromatography (RPLC) coupled to a Waters Synapt XS traveling wave-ion mobility-mass spectrometer. A 30 min RPLC gradient separation was performed with a Waters Acquity charged surface hybrid (CSH) C18 column (2.1 × 100 mm, 1.7 μm) using 60:40 acetonitrile/water with 10mM ammonium formate (Mobile Phase A) and 88:10:2 isopropanol/acetonitrile/water with 10 mM ammonium formate (Mobile Phase B) at a flow rate of 0.3 mL/min.^25^ The column was maintained at 40°C, and a 5 μL injection volume was used. The outlet of the LC system was connected directly to the electrospray ionization (ESI) source. Ionization was achieved with the following settings: capillary voltage, ±2 kV; source temperature, 150 °C; desolvation temperature, 500 °C; desolvation gas flow, 1000 L/h; cone gas flow, 50 L/h. Traveling wave ion mobility separation was performed in Nitrogen (90 mL/min flow) with a traveling wave velocity of 550 m/s and height of 40V. Data was recorded from *m/z* 50-1200 with a scan time of 0.5 s for MS1, independent MS/MS (MS^e^), and leucine enkephalin lock-mass correction functions. Separately, targeted MS/MS experiments were performed to obtain high-quality, precursor-selected fragmentation spectra. The quadrupole resolution was set to a value of 14, which provided a ±1 *m/z* isolation window. A collision energy ramp of 40-60X eV was applied in the transfer region of the instrument.

### Lipidomic Data Analysis

Raw LC-IM-MS data files were processed using Progenesis QI (v3.0, Waters). Data was lock-mass corrected using leucine enkephalin and aligned against a QC sample. Peak detection was performed, and peak areas were normalized to the total ion count as there was no significant difference in biomass based on OD readings. Lipid identification was based on accurate mass (±5 ppm) and retention time matching against lipid standards, with confirmation of headgroup and acyl tail composition by MS/MS. Skyline for small molecules was used to visualize and isolate the MS/MS acyl tail fragments and intensity distributions of isobaric lysolipids.^26^ Waters .raw files were imported into Skyline with a 0.05 Da mass tolerance and a TOF resolving power of 30,000. A fixed window of 0.6 ms was used for IM-filtering. Fatty acid intensities from product ion measurements across the duration of each respective LC method were measured and exported for analysis.

### Daptomycin Lag-Phase Extension Assays

Prior to lag-phase extension assays, bacteria were preconditioned to lipid-enriched broth overnight. Independent 2 McFarland bacterial suspensions (n=3) were used to prepare cultures in TSB that contained either DMSO or 50 μg/mL liver polar lipid extract. Following overnight growth, bacteria were pelleted by centrifugation, washed, resuspended in sterile saline, and adjusted to 0.5 McFarland. In a BSL-2 biosafety cabinet, 500 μL of bacterial suspension was mixed into 5 mL of TSB enriched with 50 mg/L of calcium chloride. Additionally, calcium enriched TSB broth was prepared with DMSO or 50 μg/mL liver polar lipid extract and daptomycin concentrations of 0, 2.5, 5, or 7.5 μg/mL. Cultures were prepared to a final volume of 200 uL in 96-well microplates using 50 μL of calcium-enriched TSB, 100 μL of TSB containing daptomycin and liver lipid extract or DMSO, and 50 μL of bacteria suspension in TSB. OD600 measurements (BioTek Epoch2, Agilent) were taken every 30 min over a 24-hr period with orbital shaking for 30 s prior to each reading. Samples were maintained at 37°C for the duration of the experiment.

## RESULTS AND DISCUSSION

### Composition of Tissue-Derived Lipid Extracts Differ in Class and Fatty Acid Content

Lipidomic analysis of the neat extracts from heart, brain, and liver tissue demonstrated the variability of phospholipid (PL) classes across the different organs (**Figure 2a**), which agreed closely with the vendor-reported distribution. Phosphatidylcholine (PC) and phosphatidylethanolamine (PE) were the most abundant phospholipid classes across all three tissue types. Liver-derived lipid extracts exhibited the highest overall abundance of phospholipids, whereas an equivalent concentration of heart lipid extract yielded the lowest intensity of phospholipids. The majority of phospholipid species contained ester linkages in both the *sn-1* and *sn-2* positions based on mass and MS/MS fragmentation spectra. However, alternative linkages were present at appreciable levels for both PCs and PEs.^27, 28^ PCs were found to have ether linked fatty acyl tails (plasmanyl-type, or PC-O), whereas PEs tended to have vinyl ether linked fatty acyl tails (plasmalogen-type, or PE-P).^27, 29^ Although difficult to confirm with the methods used here, previous studies of these lipids have indicated that the ether or vinyl ether linkages are most commonly found at the *sn-1* position of the glycerol backbone.^27, 28^ We observed that extracts derived from different tissue sources contained distinct ratios of ester-to-(vinyl)ether linked PE and PCs. Heart lipid extract contained an almost equal amount of PE and PE-P, whereas PEs were more prevalent in liver and PE-P in brain lipid extract. Conversely, only negligible amounts of PC-O were detected in the liver and brain lipid extracts while the PC-O to PC ratio was much higher in the heart lipid extract. Lipids containing only one acyl tail, or lyso-phospholipids (LysoPL), were detected at low levels in all tissue-derived lipid extracts. However, liver lipid extract contained a substantial fraction of both LysoPCs (LPC) and LysoPEs (LPE). No ether or vinyl-ether linked LysoPLs were detected in the tissue-derived lipid extracts.

**Figure 2.**
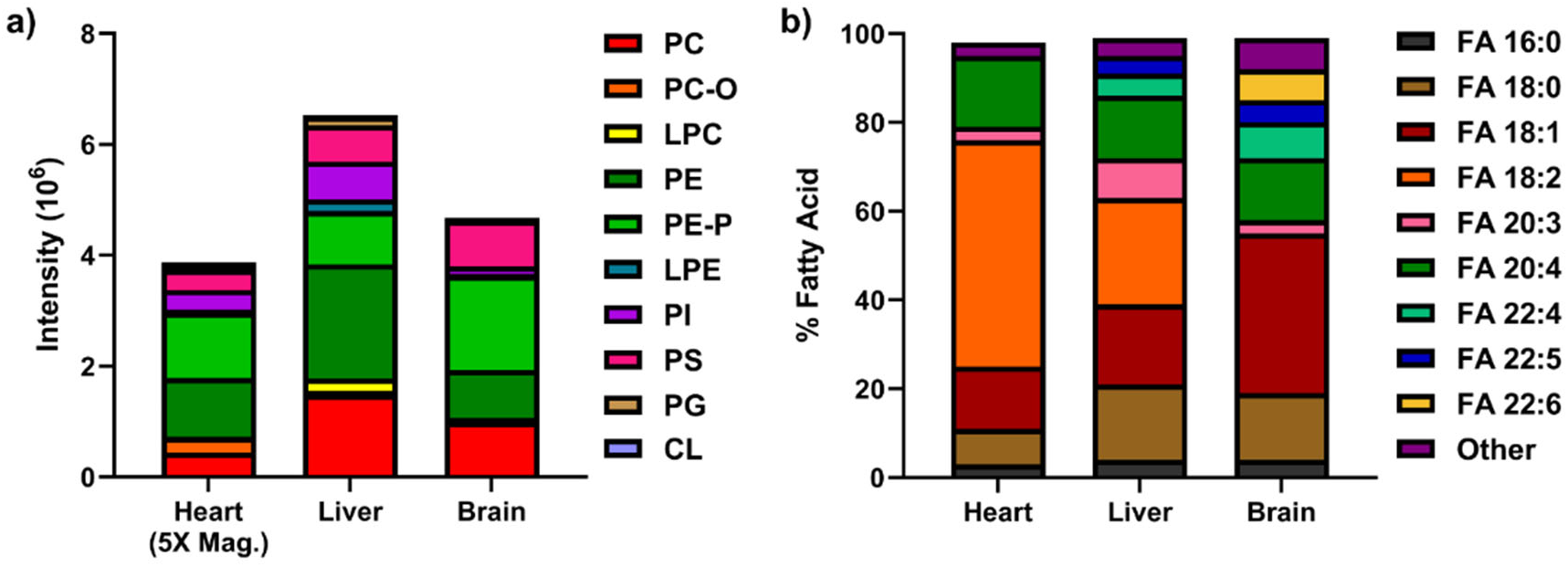
Lipidomic analysis of tissue-derived phospholipid extracts used for the supplementation of *S. aureus* cultures. a) Distribution of phospholipid classes within tissue-derived polar lipid extracts as determined by ESI-intensity. b) Distribution of fatty acyl tails within phospholipid extracts as determined by data-independent MS/MS fragmentation.

In addition to the distribution of phospholipid classes, we also evaluated the distribution of fatty acyl tails within the phospholipids derived from different tissue extracts. The fatty acyl profiles in **Figure 2b** are based on the FA fingerprints in the data-independent MS/MS spectra collected during the RPLC-IM-MS analyses. While not quantitative and susceptible to bias from fragmentation susceptibilities, the profiles nonetheless reveal substantial differences in the fatty acid content within phospholipids of heart, liver, and brain tissue extracts. The phospholipids MS/MS spectra from all tissues contained predominantly unsaturated fatty acyl tails, with minor amounts of saturated FA fragments. Heart and liver extracts contained a high proportion of FA 18:2 in their MS/MS-derived FA fingerprint, whereas the MS/MS spectra of brain lipid extract were dominated by FA 18:1.

While the proportion of FA 20:4 was similar across all extracts, the phospholipids in the brain lipid extract had high proportions of other poly-unsaturated fatty acids (PUFAs) such as FA 22:6. These analyses confirm many of the expected trends in FA and PL content from different tissue sources and demonstrate the variety of phospholipid and fatty acid structures that are available to the *S. aureus* secreted lipase, Geh.

### Growth in Lipid-Enriched Broth Yields Unsaturated S. aureus Lipids

The major phospholipid species of *S. aureus* are phosphatidylglycerol (PGs) that contain saturated and branched-chain fatty acids (BCFAs) and no endogenous unsaturated fatty acids. Under standard growth conditions, PGs 17:0/15:0 (PG 32:0) and 18:0/15:0 (PG 33:0) are usually the most abundant species but *S. aureus* PGs often contain multiple FA combinations that yield the same total carbon (*e.g.*, PG 32:0 represents mostly PG 17:0/15:0, but 18:0/14:0 and 16:0/16:0 are also detected).^5^ We first investigated whether growth of *S. aureus* in lipid-enriched broth impacted the distribution of endogenous saturated PG species. We found that the DMSO vehicle control showed no impact on native PG distributions compared to the TSB growth control **(Figure 3a)**. Growth with heart- and liver-derived lipids produced higher levels of PG 32:0 and 33:0 and minor increases of PG 34:0 (mostly 19:0/15:0) and PG 35:0 (mostly 20:0/15:0). In contrast, the addition of brain-derived lipids resulted in only a minor increase in PG 33:0 and no impact on other PG species. No changes to the fatty acyl content or distribution between branched- and straight-chain acyl tails of PGs were detected following growth of *S. aureus* in lipid-enriched broth.^25^

**Figure 3.**
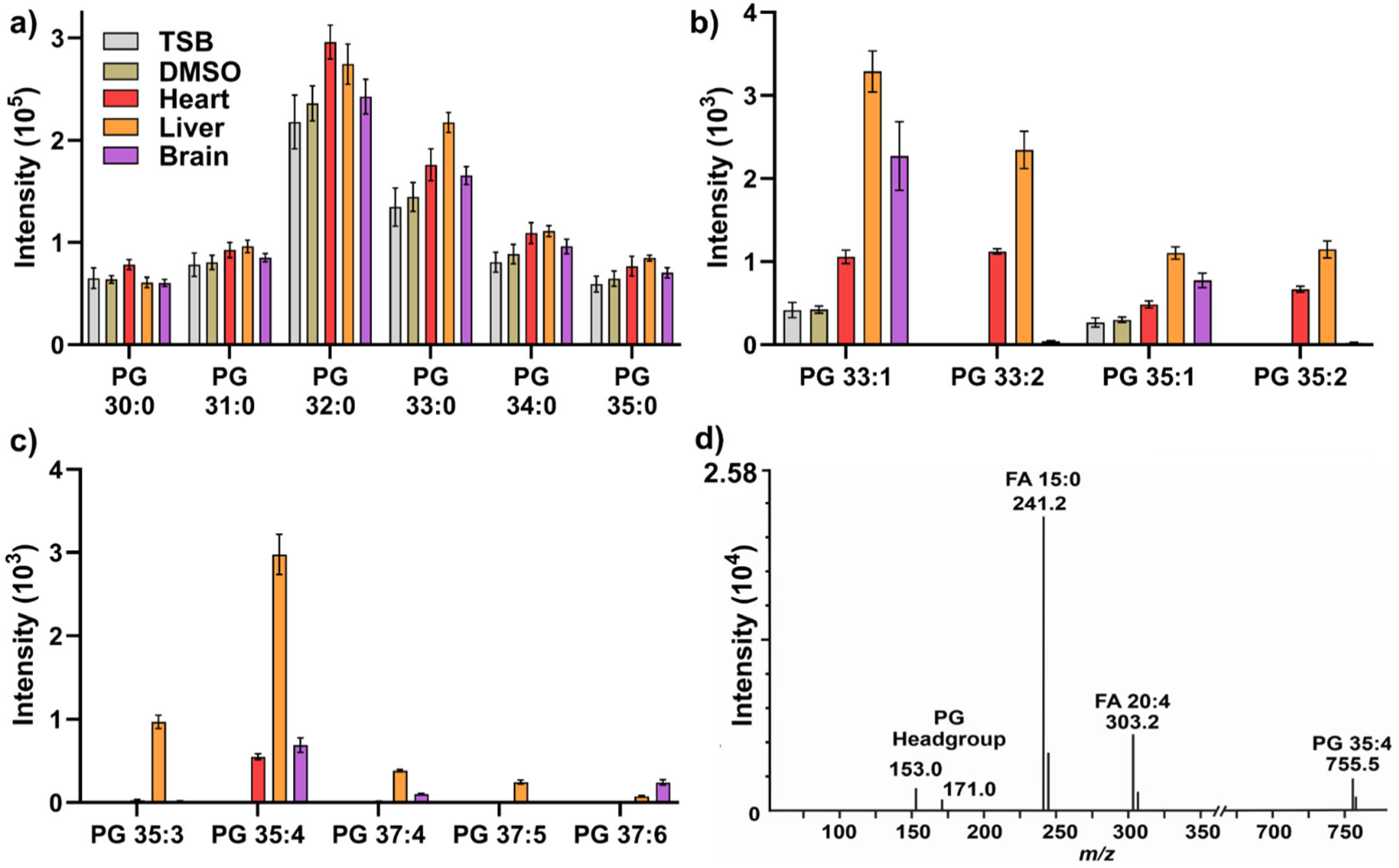
Lipidomic analysis of *S. aureus* PGs following growth in lipid-enriched TSB. Saturated (a) and unsaturated (b and c) PG profiles of *S. aureus* after overnight growth in TSB, TSB supplemented with DMSO, or TSB supplemented with 50 μg/mL of tissue-derived (heart, liver, or brain) lipid extract. d) Representative MS/MS spectra of PG 35:4 demonstrating the assignment of PG 20:4/15:0 based on relative acyl tail fragment intensities.

Despite the lack of impact on fully saturated PG species, growth of *S. aureus* in the lipid-enriched media resulted in the formation of PG species containing mono- and poly-unsaturated FAs and an odd number of total carbons (**Figure 3b**). The presence of PG species with MUFAs and PUFAs correlated with the previously reported fatty acid profiles of each tissue. Since all tissue-derived extracts contained high levels of FA 18:1 or FA 18:2, the most prominent unsaturated PGs following growth in lipid-enriched media were PG 18:1/15:0 (PG 33:1) or PG 18:2/15:0 (PG 33:2). Several PUFA-containing PG species were also detected, including PG 20:4/15:0 (PG 35:4) in all conditions and PG 22:6/15:0 (PG 37:6) in the cultures supplemented with brain-derived lipid extract (**Figure 3c**). As shown for PG 35:4 in **Figure 3d**, targeted fragmentation of these species indicated that the unsaturated FA tail was located at the *sn-1* position of the glycerol backbone, with the *sn-2* position occupied by the BCFA 15:0 as typically found in *S. aureus* lipids.^25, 30, 31^ Evidence of FASII elongation of exogenous unsaturated FAs was evident from PG species such as PGs 35:1 and 35:2 that contained FA 20:1 or FA 20:2. These FAs were not detected within the neat tissue-derived lipid extracts themselves but instead were likely elongated by two carbons from FAs 18:1 and 18:2 through FASII.^5, 14^ Note that trace amounts of PG 33:1 and PG 35:1 were detected in both TSB and DMSO controls, likely due to basal levels of FA 18:1 within TSB itself. Although incorporation of exogenous FAs into other *S. aureus* lipid classes have been reported, we did not detect any appreciable amounts of DGDGs, LysylPGs, or CLs with unsaturated fatty acids.^14, 20, 22^ Evidence of the incorporation of multiple exogenous FAs into *S. aureus* PGs was not detected either. This is likely due to the provision of intact phospholipids, which require both Geh and Fak to generate and incorporate exogenous FAs, rather than free FAs and the lower concentration of lipid used here.

### Growth in Lipid-Enriched Broth Increases S. aureus Daptomycin Tolerance

As a membrane-targeting antimicrobial, several studies in *S. aureus* and similar species have demonstrated that daptomycin tolerance is strongly influenced by the presence of unsaturated FAs in the bacterial membrane.^23, 30–33^ To determine the impact of complex lipid changes induced by growth in lipid-enriched broths, we preconditioned *S. aureus* overnight to TSB supplemented with liver-derived lipid extract and then monitored the growth of *S. aureus* in the presence of TSB supplemented with lipids and daptomycin **(Figure 4)**. The addition of liver polar lipid extract, which resulted in the highest levels of unsaturated PGs, had little influence on the growth of *S. aureus* compared to the DMSO vehicle control, nor did the addition of daptomycin at 2.5 μg/mL. Delayed entrance into log-phase growth was observed for both lipid-supplemented and control cultures at a daptomycin concentration of 5 μg/mL, with the lipid-supplemented culture reaching mid-log phase approximately 30 min earlier than the DMSO control. Severe delays in growth were induced by 7.5 μg/mL daptomycin for both culture conditions. However, *S. aureus* grown in lipid-enriched TSB achieved some growth after 12 hrs whereas the DMSO control condition failed to grow until 18+ hrs. These results demonstrate the membrane lipid composition of *S. aureus in vivo*, which is expected to contain host-derived unsaturated FAs, promotes tolerance to membrane-targeted daptomycin. As standard laboratory culture conditions fail to replicate the *in vivo* membrane composition of *S. aureus*, there is a risk that standard antimicrobial susceptibility testing practice underestimates the organism’s tolerance of daptomycin after exposure to the lipid-rich infection environment of mammalian tissues.

**Figure 4.**
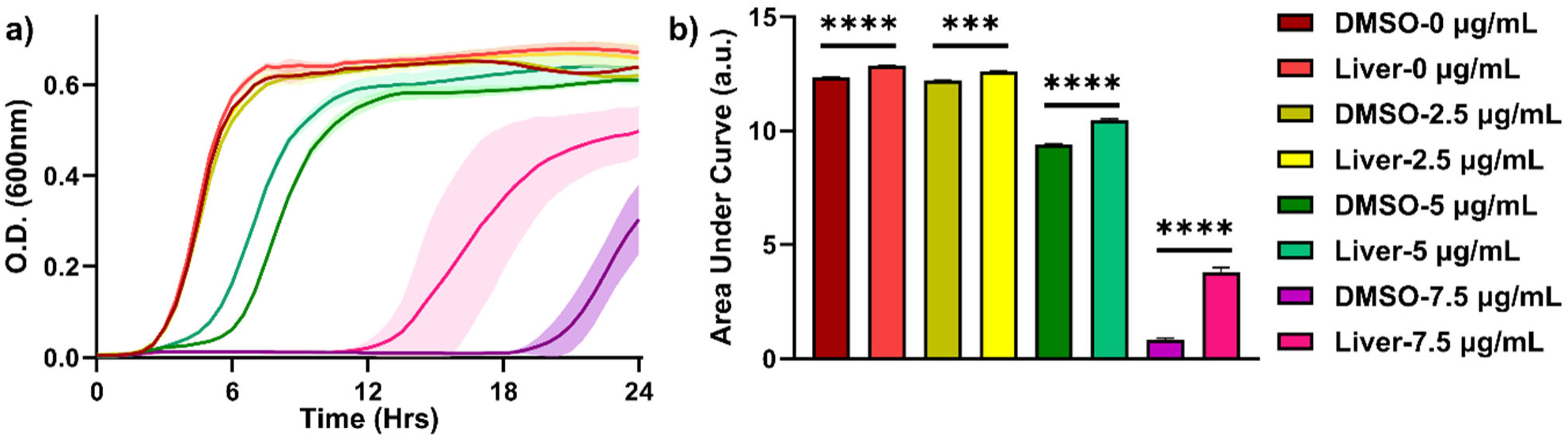
Influence of liver-derived lipid supplementation on the growth of *S. aureus* with daptomycin. Growth curves (a) and area under the curve (b) for *S. aureus* JE2 (n=3) cultured in Ca^2+^ modified TSB containing DMSO or 50 µg/mL liver polar lipid extract and 0, 2.5, 5.0, or 7.5 µg/mL of daptomycin. *S. aureus* was pre-conditioned overnight with DMSO or 50 µg/mL liver polar lipid extract prior to monitoring growth curves. *P*-values were calculated using multiple unpaired t-tests, where ***<0.005 and **** < 0.00005.

### Impact of S. aureus Lipase Activity on Extracellular Phospholipids

While the influence of exogenous FAs on the membrane lipids of *S. aureus* is well studied, comparatively little investigation has been done into the influence that *S. aureus* lipase activity has on the phospholipid species in the extracellular domain. We therefore carried out a comparative analysis of the lipid-enriched TSB before and after it was exposed to *S. aureus* to identify the phospholipid classes and species that were most significantly impacted by secreted lipase activity. Our results revealed that the type of glycerol linkage and phospholipid headgroup influenced the extent to which the extracellular phospholipids were degraded after growth with *S. aureus* **(Figure 5a)**. As expected, phospholipid species containing ether or vinyl-ether linkages to the glycerol backbone, such as PC-O and PE-P, were largely unaffected (*ca.* 60-80% remaining) by growth with *S. aureus* as the ether bonds are not susceptible to hydrolysis by Geh. It is likely that the small reduction in PC-O and PE- P species is attributed to hydrolysis of the acyl tail at the *sn-2* position, which is less favorable for Geh but still feasible.^18^ Within ester-linked phospholipids, we observed that PEs were the most impacted of the major phospholipid classes present within the lipid-enriched broths. The lipidomics data revealed that the PE intensity in the spent lipid-enriched TSB was only 5% of the PE intensity detected in the lipid-enriched broth that was not exposed to *S. aureus*, corresponding to a 95% reduction in PEs. Although other phospholipid classes were similarly affected, the amount of these lipids in the tissue-derived extracts was significantly lower to start with **(Figure 2a)**. These results demonstrate that the prominent *S. aureus* lipase, Geh, acts on acyl tails of phospholipid classes having anionic (PG), acidic (PS), and neutral (PI) headgroups.

**Figure 5.**
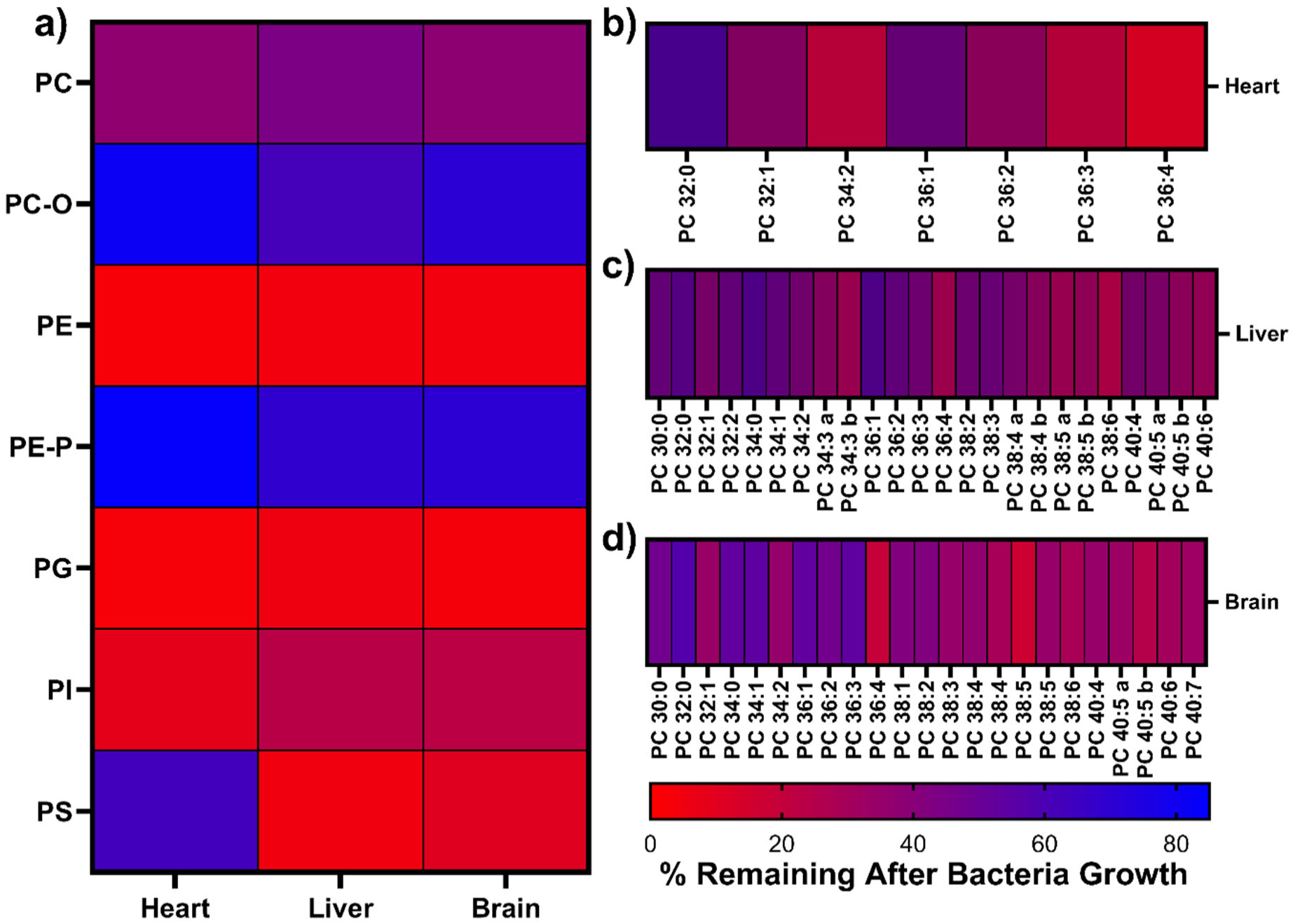
The influence of *S. aureus* on extracellular phospholipid composition. a) Heat map showing the percent of phospholipids remaining in spent lipid-enriched broth following growth of *S. aureus*. The impact of *S. aureus* growth on specific PC species for spent TSB containing b) hearth, c) liver, and d) brain lipid extracts. Lipid species with multiple RPLC peaks are distinguished by appending the lipid name with *“a”* and *“b*.” The percent remaining was determined relative to fresh lipid-supplemented TSB was exposed to the same conditions but without the addition of *S. aureus*.

To our surprise, PCs in lipid-enriched broths were less impacted following growth with *S. aureus* than other phospholipid species. The intensity of PCs was approximately 60% lower in the spent broth relative to the lipid-enrich broth that was not exposed to *S. aureus*. PCs were one of the most abundant phospholipid species in the tissue-derived lipid extracts, representing a significant fraction of accessible ester-linked acyl tails. We further investigated the impact of *S. aureus* exposure on individual PC species to determine if this intermediate level of degradation was the result of heterogeneous lipase activity within the lipid class **(Figure 5b-d)**. Although individual tissues had different PC species available, trends related to the length and degree of unsaturation of the acyl tails emerged. The most evident trend was that PCs with the same chain length but higher degrees of unsaturation were hydrolyzed to a greater extent. This is most clear in the heart-derived PCs with 36:1 to 36:4 acyl tail compositions, where the percent remaining steadily decreased as the degree of unsaturation increased. The longer and highly-unsaturated PCs in liver- and brain-derived lipid extracts showed the same trend, but little difference was detected beyond 5 degrees of unsaturation. Finally, there were modest differences in the amount of isobaric PC species (*i.e.*, same *m/z* but different retention time) remaining after *S. aureus* growth. This suggests that subtle differences in the structure of the acyl tails could influence the lipid’s susceptibility to lipase activity. While the detailed kinetics of acyl tail hydrolysis are outside the scope of this manuscript, these data suggest that structural factors such as head group polarity, glycerol backbone linkage, and acyl tail length and degree of unsaturation influence *S. aureus* lipase activity and thereby the extracellular lipid content.

The activity of *S. aureus* secreted lipase was also investigated from a second perspective by evaluating the products of glycerol ester hydrolysis of phospholipids. As shown in **Figure 1**, cleavage of an acyl tail from a phospholipid yields a free fatty acid and a lysophospholipid (LysoPL or LPL) in which the remaining acyl tail can be located at the *sn-1* (1-acyl lysolipids) or *sn-2* (2-acyl lysolipid) position of the backbone. The lipidomics data collected for lipid-enriched broths before and after exposure to *S. aureus* were evaluated for the presence of LysoPL species **(Figure 6)**. As shown in **Figure 6a**, the intensity of LysoPL increased in broths supplemented with heart, liver, and brain extract following overnight growth of *S. aureus*. The distribution of LysoPL classes mirrored that of the initial phospholipid composition of the tissue-derived extracts with the highest intensity from LysoPCs (LPCs) and LysoPEs (LPEs). Some tissue sources, particularly liver, natively contained a large proportion of LPCs and LPEs, which were elevated further following exposure to *S. aureus* and supplemented by other species of LysoPLs such as LPI, LPG, and LPS. Given the reported preference of Geh for the *sn-1* position, we evaluated the distribution of 1-acyl vs. 2-acyl LysoPLs both before and after the lipid-enriched broth was exposed to *S. aureus*.^18^ Based on the chromatographic separation of LysoPL isomers, we observed that the fresh lipid-enriched broths (and the tissue-derived lipid extracts themselves) contained predominately the 1- acyl type LysoPLs **(Figure 6b)**. While 1-acyl LysoPLs are thought to be the more prevalent isomers in mammalian tissues, such interpretations are difficult to make based purely on lipidomics measurements as the instability of 2-acyl LysoPLs can result in spontaneous migration of the acyl tail to form the more stable 1-acyl LysoPLs.^18, 34–36^ Nonetheless, we detected a significant increase in the intensity of 2-acyl LysoPLs in the spent lipid-enriched broths. This resulted in a shift in the ratio of LysoPL isomers from 70-90% 1-acyl in the fresh broth to 45-55% 2-acyl following *S. aureus* growth.

**Figure 6.**
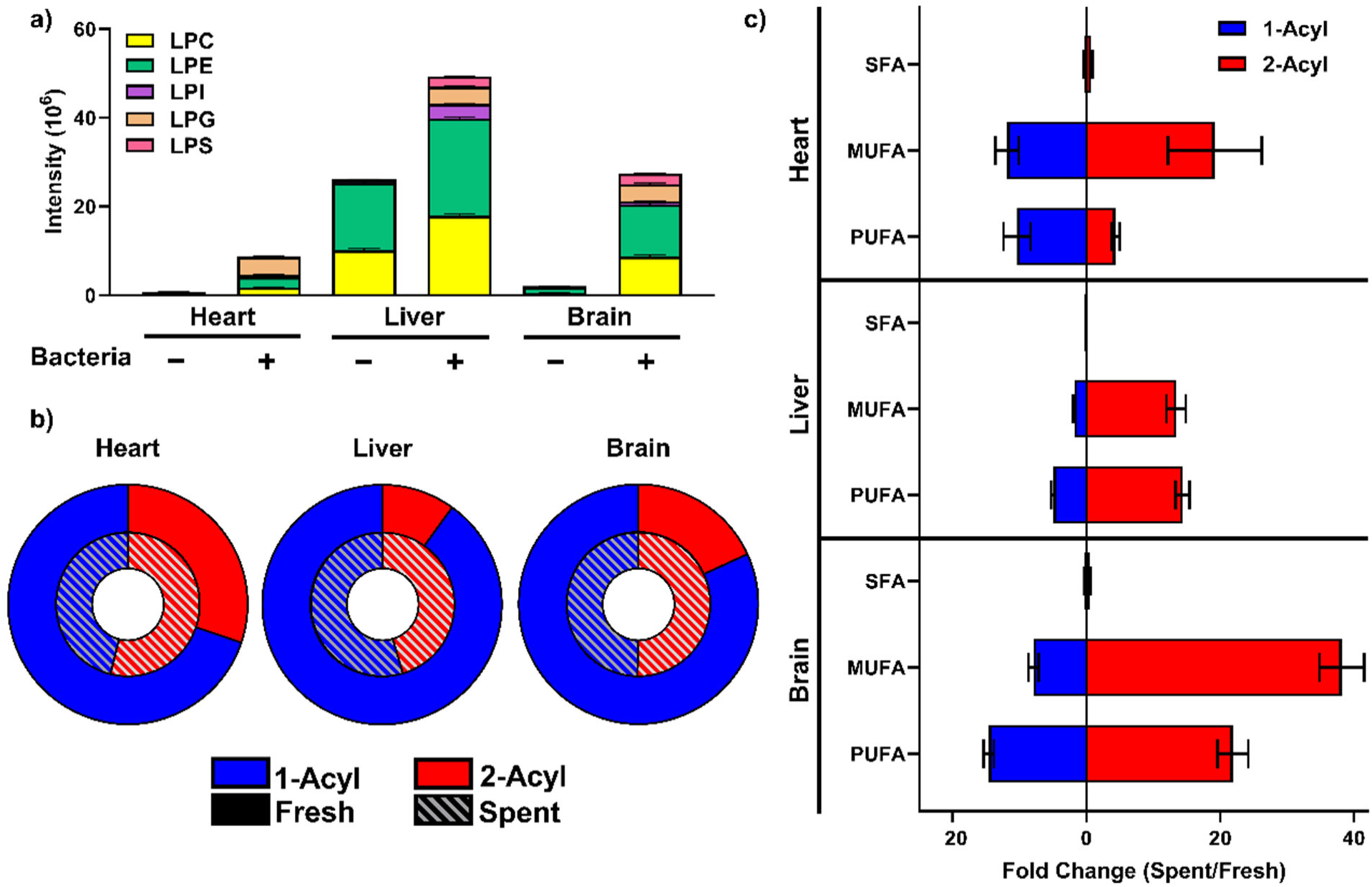
Alteration of lysophospholipid levels and structures following *S. aureus* growth. a) Total summed abundance of lysophospholipid classes in fresh (Bacteria -) and spent (Bacteria +) lipid-enriched TSB. b) Distribution of 1-acyl vs. 2-acyl lysophospholipids in fresh and spent lipid-enriched TSB. c) Fold change differences in saturated (SFA), monounsaturated (MUFA) and polyunsaturated (MUFA) 1-acyl and 2-acyl lysophospholipid species between spent and fresh lipid-enriched broth.

The changes in LysoPL levels were not uniform across all species, but instead varied based on the acyl tail structure. As shown in **Figure 6c**, LysoPLs containing saturated acyl tails were reduced after culture with *S. aureus* for all the lipid-enriched broths evaluated (*i.e.*, fold-change <1). This change was especially significant in the liver extract-enrich broth where a large proportion of saturated LysoPCs and LysoPEs were present in the supplemented broth at the start. Although the distribution in 1-acyl to 2-acyl LysoPLs shifted, the general increase in total LysoPL species was driven by higher amounts of both the 1-acyl and 2-acyl forms of LysoPLs with mono- and poly-unsaturated acyl tails. However, the fold-change increase of 2-acyl LysoPLs between spent and fresh lipid-enriched broths was greater for all conditions except PUFA-containing LysoPLs in TSB supplemented with heart-derived lipid extract. This is consistent with the general preference within mammalian phospholipids for the *sn-2* position to contain unsaturated fatty acyl tails, whereas saturated FAs are more often found at the *sn-1* position. The net increase in LysoPLs species following growth of *S. aureus* in lipid-enriched broth has important implications when translated to the infection environment, including the modulation of immune and inflammatory responses. LysoPCs are well-known lipid mediators of pro- and anti-inflammatory responses, where the distinction between the promotion or suppression of inflammation is determined by the acyl tail.^34^ For example, saturated and monounsaturated LPCs such as 16:0, 18:0, and 18:1 are known to stimulate cytokine secretion to promote inflammatory response.^37^ On the other hand, polyunsaturated species like LPC 20:4 and 22:6 are strongly anti-inflammatory.^38^ LysoPEs are associated with several signaling pathways and aberrantly high levels of LysoPEs are implicated in blood clot formation, damage to endothelial cells, as well as neuronal death.^39, 40^ Furthermore, receptors of LysoPLs are able to distinguish 1-acyl and 2-acyl forms and often show preference for 2-acyl species, especially those containing unsaturated FA species.^41, 42^ Given the important physiological roles of LysoPLs in mammals, there is strong potential that the formation of LysoPLs through the activity of *S. aureus* secreted lipases influences the host inflammatory response and other processes at the site of infection.

## CONCLUSIONS

Our findings demonstrate that *S. aureus* can hydrolyze exogenous ester-linked phospholipids from a variety of phospholipid classes that are found within the lipid extracts from different mammalian tissues. The resulting free fatty acids, consisting primarily of mono- and poly-unsaturated species, are then incorporated into the major phospholipid of the *S. aureus* membrane, phosphatidylglycerol, at the *sn-1* position of the glycerol backbone. Our data revealed that the specific distribution of unsaturated PG species resulting from these processes varied across the different sources of lipid extract used to enrich the broth, such that the unsaturated PGs largely mirrored the FA composition of the extract. Although most poly-unsaturated FA species are inherently toxic to *S. aureus*, their incorporation into its membrane lipids is likely a detoxification mechanism while simultaneously abstracting the PUFAs from the extracellular environment where they may contribute to inflammatory signaling.^8, 12, 32, 43^ On the other hand, the addition of highly-unsaturated FAs into the acyl tails of *S. aureus* PGs has a strong fluidizing influence on the membrane that may be beneficial to its survival in the host environment and tolerance against membrane-targeting antimicrobials.^20, 33^ This conclusion is supported by our evidence that *S. aureus* pre-conditioned to and cultured in broth containing liver-derived lipids is better able to grow in the presence of high concentrations of daptomycin.

In the context of the extracellular environment, the degradation of phospholipids by secreted *S. aureus* lipase dramatically increases the native pool lysophospholipid species and shifts the distribution from the typical 1-acyl form toward the formation of 2-acyl species. These results are consistent with evidence that Geh acts similarly to Phospholipase A_1_ with a preference for the acyl tails at the *sn-1* position of the phospholipid glycerol backbone.^18^ However, our results extend these findings to demonstrate that *sn-1* preference applies to many phospholipid classes, including PEs and PCs that are highly-abundant in mammalian cells and tissues. At the same time, the general rise in LysoPLs and the specific increase in 2-acyl species in the extracellular environment holds significant implications for the inflammatory and immune response of the host to *S. aureus* infection. The binding of LysoPLs by GPCRs serves as the activating or inhibitory signal for a variety of cellular responses ranging from inflammation to calcium homeostasis^39–42^. While their specific role in response to *S. aureus* infection has not been investigated, the potential that lipase-generated LysoPLs contribute to the host immune response is worthy of future exploration. Collectively, the results of this work demonstrate that the lipidome of both the bacterium and the host are altered by the action of staphylococcal lipase activity, with implications from antibiotic susceptibility to immune response.

## ACKNOWLEDGEMENTS

Funding was provided by the National Institute of Allergy and Infectious Disease to K.M.H. (R01AI173144).

## SUPPORTING INFORMATION

Excel File (.xlsx) - Acyl tail compositions determined by targeted MS/MS of bacteria pellets, fresh and spent lipid-enriched broths.

## DATA AVAILABILITY

The raw RPLC-IM-MS and RPLC-IM-MS/MS data files from the experiments performed for the present study are available through the MassIVE repository under accession number MSV000102454.

## Notes

### Competing Interest Statement

The authors have declared no competing interest.

